# A non-classical mechanism of β-lactam resistance in Methicillin-Resistant *Staphylococcus aureus* (MRSA) and its effect on virulence

**DOI:** 10.1101/2022.06.17.496612

**Authors:** Nidhi Satishkumar, Li-Yin Lai, Nagaraja Mukkayyan, Bruce E. Vogel, Som S. Chatterjee

**Author notes:** Corresponding Author: Som S. Chatterjee, Columbus Center, 701 Pratt Street, Baltimore, MD 21202.

## Abstract

Methicillin-Resistant *Staphylococcus aureus* (MRSA) are pathogenic bacteria that are infamously resistant to β-lactam antibiotics, a property attributed to the *mecA* gene. Recent studies have reported that mutations associated with the promoter region of *pbp4* demonstrated high levels of β-lactam resistance, suggesting the role of PBP4 as an important non-*mecA* mediator of β-lactam resistance. The *pbp4* promoter-associated mutations have been detected in strains with or without *mecA*. Our previous studies that were carried out in strains devoid of *mecA* described that *pbp4* promoter-associated mutations lead to PBP4 overexpression and β-lactam resistance. In this study, by introducing various *pbp4* promoter-associated mutations in the genome of an MRSA strain, we demonstrate that PBP4 overexpression can supplement *mecA*-associated resistance in *S. aureus* and can lead to increased β-lactam resistance. The promoter and regulatory region of *pbp4* is shared with a divergently transcribed gene, *abcA*, which encodes for a multidrug exporter. We demonstrate that the promoter mutations caused an upregulation of *pbp4* and downregulation of *abcA*, confirming that the resistant phenotype is associated with PBP4 overexpression only. PBP4 has also been associated with staphylococcal pathogenesis, however, its exact role remains unclear. Using a *C. elegans* model, we demonstrate that strains having increased PBP4 expression are less virulent compared to wild-type strains, suggesting that β-lactam resistance mediated via PBP4 likely comes at the cost of virulence.

**Importance:** Our study demonstrates the ability of PBP4 to be an important mediator of β-lactam resistance in not only Methicillin-susceptible *Staphylococcus aureus* (MSSA) background strains as previously demonstrated, but also in MRSA strains. When present together, PBP2a and PBP4 overexpression can produce increased levels of β-lactam resistance, causing complications in treatment. Thus, this study suggests the importance of monitoring PBP4-associated resistance in clinical settings, as well as understanding the mechanistic basis of associated resistance, so that treatments targeting PBP4 may be developed. This study also demonstrates that *S. aureus* strains with increased PBP4 expression are less pathogenic, providing important hints about the role of PBP4 in *S. aureus* resistance and pathogenesis.

## Introduction

*Staphylococcus aureus* is a Gram-positive pathogen that can cause skin and soft tissue infections (SSTIs), bacteremia, osteomyelitis, and sepsis in humans (1). Along with being equipped with a wide array of virulence factors, *S. aureus* is also resistant to a wide range of antibiotics, making infections difficult to treat (2). In particular, Methicillin-Resistant *Staphylococcus aureus* (MRSA) is infamous for being resistant to β-lactams, resulting in over 120,000 deaths in 2019 globally (3). β-lactams are a class of antibiotics known for their safety, efficacy and tissue distribution which makes them the most commonly prescribed antibiotics (4). β-lactams bind to Penicillin Binding Proteins (PBPs), which are integral proteins involved in the final stages of cell wall synthesis. The binding of β-lactams to PBPs causes their inactivation, leading to weakening of the cell wall and subsequently, cell death (5). MRSA contains the gene *mecA*, which encodes PBP2a, a PBP that has decreased affinity towards β-lactams, allowing cells to survive even in high concentrations of β-lactams (2).

Historically, resistance mechanisms in *S. aureus* have been acquired in waves. With the introduction of every new generation of β-lactams, *S. aureus* has been able to develop new resistance mechanisms (2). This ability of *S. aureus* to constantly develop resistance mechanisms makes it important to focus on other, novel mechanisms of resistance. Keeping this in mind, our previous studies involved serial passaging of strains in increasing amounts of β-lactams with the aim of identifying non-*mecA* mechanisms of resistance (6). We determined that mutations associated with the promoter region of PBP4 were largely prevalent in resistant strains (6, 7). Various studies by other groups also identified *pbp4* associated mutations in laboratory-generated (8, 9) as well as in clinically isolated (10, 11) resistant strains of both, MRSA and MSSA backgrounds, suggesting the clinical relevance of these mutations. PBP4, a non-essential, low molecular weight PBP in *S. aureus*, is produced in low amounts, and its role in resistance or pathogenesis is not very well described (12). We previously demonstrated that *pbp4* promoter-associated mutations led to significant β-lactam resistance in strains devoid of *mecA* by causing increased PBP4 expression and enhanced cell wall cross-linking (7). However, the effect of these mutations in strains containing *mecA* is unknown. Thus, in this study, we inspected whether PBP4, through the means of overexpression, supplemented *mecA*-mediated β-lactam resistance. We used the strain SF8300, a derivative of USA300, which is one of the most prominent community-associated MRSA (CA-MRSA) strains detected in the U.S.(2). *pbp4* promoter-associated mutations that were detected in a laboratory passaged, resistant strain that was devoid of *mecA* (namely CRB) and in a resistant strain containing *mecA* (namely CRT) were introduced in SF8300 (6). Further, a *pbp4* promoter-associated mutation detected in a β-lactam resistant, clinical isolate (namely Strain 1) was also used in this study (13). Using these promoter mutants, we demonstrated that *pbp4* promoter-associated mutations leads to PBP4 overexpression and increased β-lactam resistance in MRSA strains in a mechanism similar to what was previously shown in MSSA strains (7, 13).

*pbp4* shares its promoter and regulatory region with *abcA*, a gene that encodes for an ATP-binding cassette transporter protein. ABC transporters are notorious for enabling resistance to chemotherapeutic agents in both prokaryotes and eukaryotes via export activity. Since AbcA has previously been associated with antibiotic resistance (14), we were interested in understanding whether the promoter-associated mutations altered *abcA* expression that potentially led to AbcA-mediated β-lactam resistance. However, our results indicated that while the promoter-associated mutations led to increased *pbp4* expression, they resulted in downregulation of *abcA*, indicating that AbcA did not have any role in β-lactam resistance and that the resistance phenotype observed in the promoter mutants was solely attributed to PBP4.

Along with β-lactam resistance, PBP4 has recently been associated with virulence in previous studies. However, the exact role of PBP4 in virulence remains unclear, as these studies had contrasting reports (15, 16). In order to get a better understanding of the role of PBP4 in virulence, we used a *C. elegans* infection model and demonstrated that MRSA strains containing *pbp4* promoter-associated mutations had decreased virulence compared to the wild-type strain.

Our findings suggest the importance of monitoring PBP4-associated resistance in both MRSA and MSSA strains and indicate that treatment options would potentially have to consider targeting both PBP2a and PBP4, as when present, both the proteins independently mediate β-lactam resistance via distinct mechanisms leading to increased resistance. Our results also confirm that promoter-associated mutations only allow for PBP4 overexpression and do not facilitate AbcA-mediated resistance, indicating that PBP4-associated resistance has the potential of being a prominent resistance mechanism in the future. Finally, our results also provide important clues associated with the role of PBP4 in virulence and suggest that strains with increased PBP4 may have decreased virulence, a phenomenon that needs to be studied further in details in the future.

## Results

### *pbp4* promoter-associated mutations led to increased PBP4 expression and β-lactam resistance in MRSA strains

Our previous studies demonstrated that *pbp4* promoter-associated mutations led to PBP4 overexpression that subsequently resulted in β-lactam resistance (7, 13). These studies were performed by introducing promoter-associated mutations into strains devoid of *mecA*. Using allelic replacement, we created isogenic mutants by introducing three different, previously detected mutations in the *pbp4* promoter region of SF8300 (13, 17). Of these, two mutations were detected in laboratory-generated resistant strains as a result of a passaging experiment. The first mutation was detected in the strain CRB, and was a 36 bp duplication 290 bps upstream of the *pbp4* start codon. This mutation was introduced into the *pbp4*-promoter region of SF8300, giving rise to the strain SF8300 P*pbp4*\* (CRB). Similarly, insertion of mutations detected in the strain CRT (a thymine insertion 377 bp upstream the start codon and a 90 bp deletion 275 bp upstream the start codon) into the *pbp4*-promoter region of SF8300 produced SF8300 P*pbp4*\* (CRT). The third mutation was one detected in a clinically-isolated strain (10, 13), namely Strain 1, and consisted of a T to A substitution 266 bp upstream the start codon. The introduction of this mutation in SF8300 gave rise to SF8300 P*pbp4*\* (Strain 1) **(Figure 1a)**.

**Figure 1:**
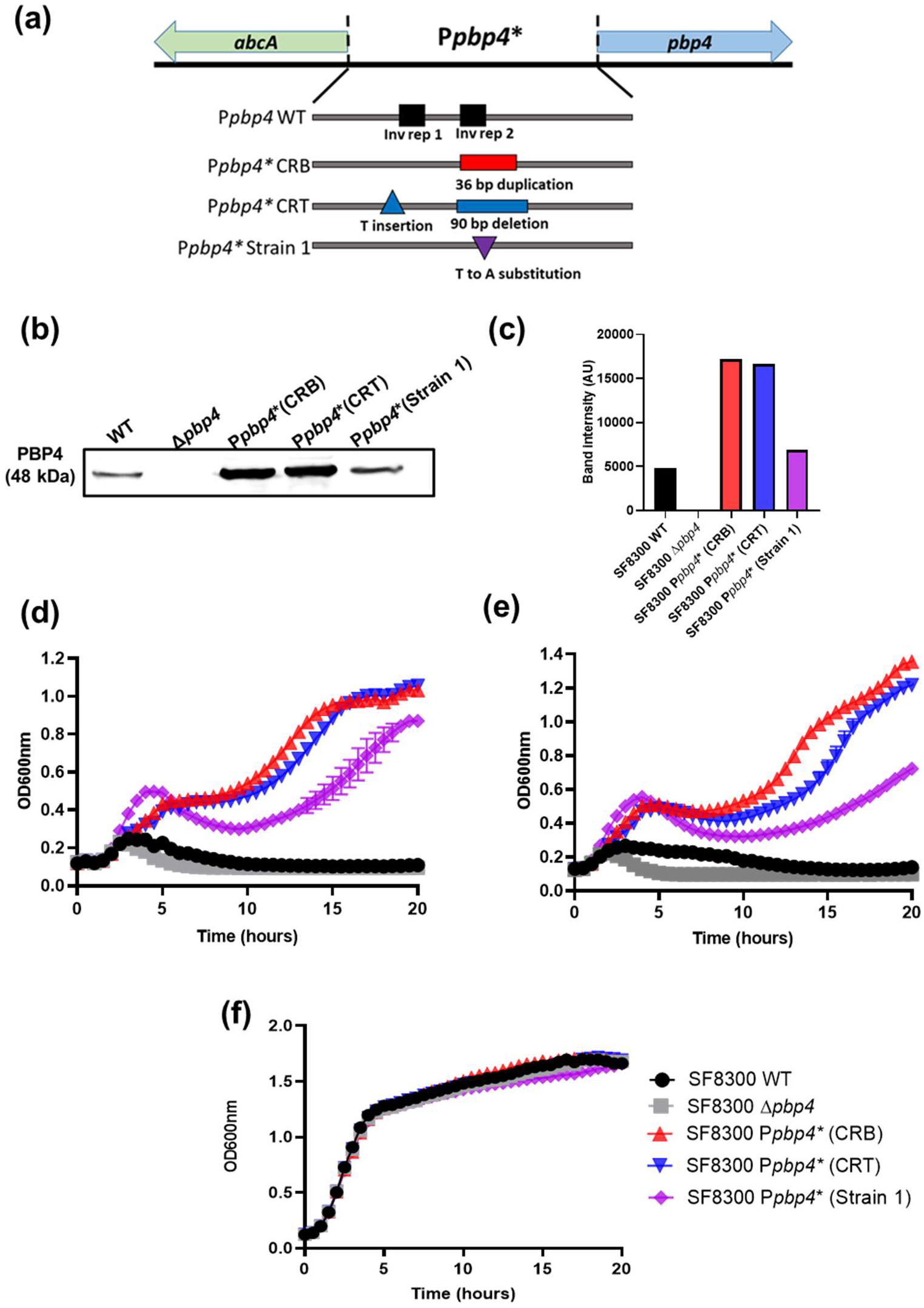
*pbp4* promoter-associated mutations caused PBP4 overexpression and increased β-lactam resistance in MRSA strains. **(a) Schematic diagram of the *pbp4*-P*pbp4*-*abcA* region**. The *pbp4* and *abcA* transcriptional start sites are separated by 420 bps of the promoter region. Along with the wild-type promoter (P*pbp4* WT), promoter mutations were seen in the strain CRB (36 bp duplication at 290 bp upstream the start codon), CRT (T insertion at 377 bp and a 90 bp deletion at 275 bp upstream the start codon) and Strain 1 (T to A substitution 266 bp upstream the start codon) are represented. **(b) Immunoblotting of PBP4 expression levels among selected strains**. Proteins from the membrane fraction of the WT strain (SF8300) and strains containing promoter-associated mutations (SF8300 P*pbp4*\* (CRB), SF8300 P*pbp4*\* (CRT) and SF8300 P*pbp4*\* (Strain 1) were probed with an antibody specific to PBP4 and were visualized for protein expression. **(c)Densitometry analysis of PBP4 immunoblotting**. Compared to WT, the strains with promoter mutations had increased levels of PBP4 (48 kDa). Δ*pbp4* was used as a control, where there was no PBP4 band detected. **(d) Growth assay of SF8300 WT and mutant strains in the presence of 4 µg/mL Nafcillin**. Strains containing promoter-associated mutations showed significantly enhanced survival compared to the WT and Δ*pbp4* strains. (SF8300 WT versus SF8300 P*pbp4*\* (CRB), SF8300 WT versus SF8300 P*pbp4*\* (CRT) and SF8300 WT versus SF8300 P*pbp4*\* (Strain 1), P-value <0.0001) **(e) Growth assay of SF8300 WT and mutant strains in the presence of 8 µg/mL Oxacillin**. Strains containing promoter-associated mutations showed significantly enhanced survival compared to the WT and Δ*pbp4* strains. (SF8300 WT versus SF8300 P*pbp4*\* (CRB), SF8300 WT versus SF8300 P*pbp4*\* (CRT) and SF8300 WT versus SF8300 P*pbp4*\* (Strain 1), P-value <0.0001) **(f) Growth assay of SF8300 WT and mutant strains in the absence of antibiotics**. There was no significant difference in growth pattern amongst the selected isogenic strains in absence of antibiotics.

In order to determine if introduction of these mutations affected PBP4 expression in the selected isogenic strains, immunoblotting was performed using an antibody specific to PBP4. Compared to the wild-type (WT) strain SF8300, strains containing promoter-associated mutations had significantly increased expression of PBP4 **(Figure 1b)**. SF8300 P*pbp4*\* (CRB) had the highest amount of expressed protein, followed by SF8300 P*pbp4*\* (CRT) and SF8300 P*pbp4*\* (Strain 1). Δ*pbp4* was used as a control, for which there was no PBP4 band detected **(Figure 1c)**. These results indicated that PBP4 overexpression occurred as a result of promoter-associated mutations in MRSA strains, in a manner similar to what was previously detected in MSSA strains (7, 13).

Following confirmation of PBP4 overexpression, we performed a growth assay in presence of β-lactams such as nafcillin and oxacillin to examine the effect of PBP4 overexpression on β-lactam resistance. When exposed to β-lactams, strains with promoter-associated mutations were able to survive significantly better in presence of either of the antibiotics, when compared to the WT and Δ*pbp4* strains for 4 µg/mL nafcillin **(Figure 1d)** and 8 µg/mL oxacillin **(Figure 1e)**. The wild-type strain as well as the strains with promoter-associated mutations all had similar growth patterns in absence of β-lactams **(Figure 1f)**. Taken together, these results suggested that promoter-associated mutations led to PBP4 overexpression and β-lactam resistance in MRSA background, similar to what was previously demonstrated in MSSA background strains (7, 13).

### pbp4 promoter-associated mutations led to increased expression of pbp4 and decreased expression of abcA

The *pbp4* gene shares its 420 bp promoter and regulatory region with a neighboring, divergently transcribed gene, namely *abcA* (18). AbcA is an ATP Binding Cassette-like transporter protein that has been reported to export various chemicals, dyes and antibiotics, thus contributing to antibiotic resistance in *S. aureus* (14, 19). It also has the ability to export Phenol Soluble Modulins (PSMs) that are cytolytic toxins, thus also plays a role in *S. aureus* virulence (20). Since the mutations detected upstream of *pbp4* start codon also lie upstream of the *abcA* start codon **(Figure 1a)**, we were interested in whether they caused any alterations in the expression of *abcA* that subsequently contributed to β-lactam resistance in *S. aureus*. We thus performed qRTPCR to examine the expression pattern of *pbp4* and *abcA* in presence of promoter-associated mutations. At 4 hours, *pbp4* transcripts were expressed in very low amounts in SF8300 WT, and were significantly increased in strains containing promoter-associated mutations **(Figure 2a;** SF8300 P*pbp4* (WT) versus SF8300 P*pbp4*\* (CRB), P-value < 0.0001, SF8300 WT versus SF8300 P*pbp4*\* (CRT), P-value < 0.0001, SF8300 WT versus SF8300 P*pbp4*\* (Strain 1), P-value = 0.0004). On the other hand, *abcA* transcripts had increased expression in SF8300 WT but had significantly decreased expression in strains containing promoter-associated mutations **(Figure 2b;** SF8300 WT versus SF8300 P*pbp4*\* (CRB), P-value = 0.0161, SF8300 WT versus SF8300 P*pbp4*\* (CRT), P-value = 0.0016, SF8300 WT versus SF8300 P*pbp4*\* (Strain 1), P-value = 0.0203). These results indicate that while the promoter-associated mutations caused overexpression of *pbp4*, but led to a downregulation of *abcA*. Thus, the increased β-lactam resistance seen due to the promoter-associated mutations was solely attributed to PBP4 overexpression and not AbcA activity. The results of qRTPCR also corresponded with the results seen with PBP4 immunoblotting in terms of different levels of PBP4 overexpression, which indicated that SF8300 P*pbp4*\* (CRB) had the highest amount of PBP4 overexpression, followed by SF8300 P*pbp4*\* (CRT) and SF8300 P*pbp4*\* (Strain 1) **(Figure 2a, Figure 1c)**.

**Figure 2:**
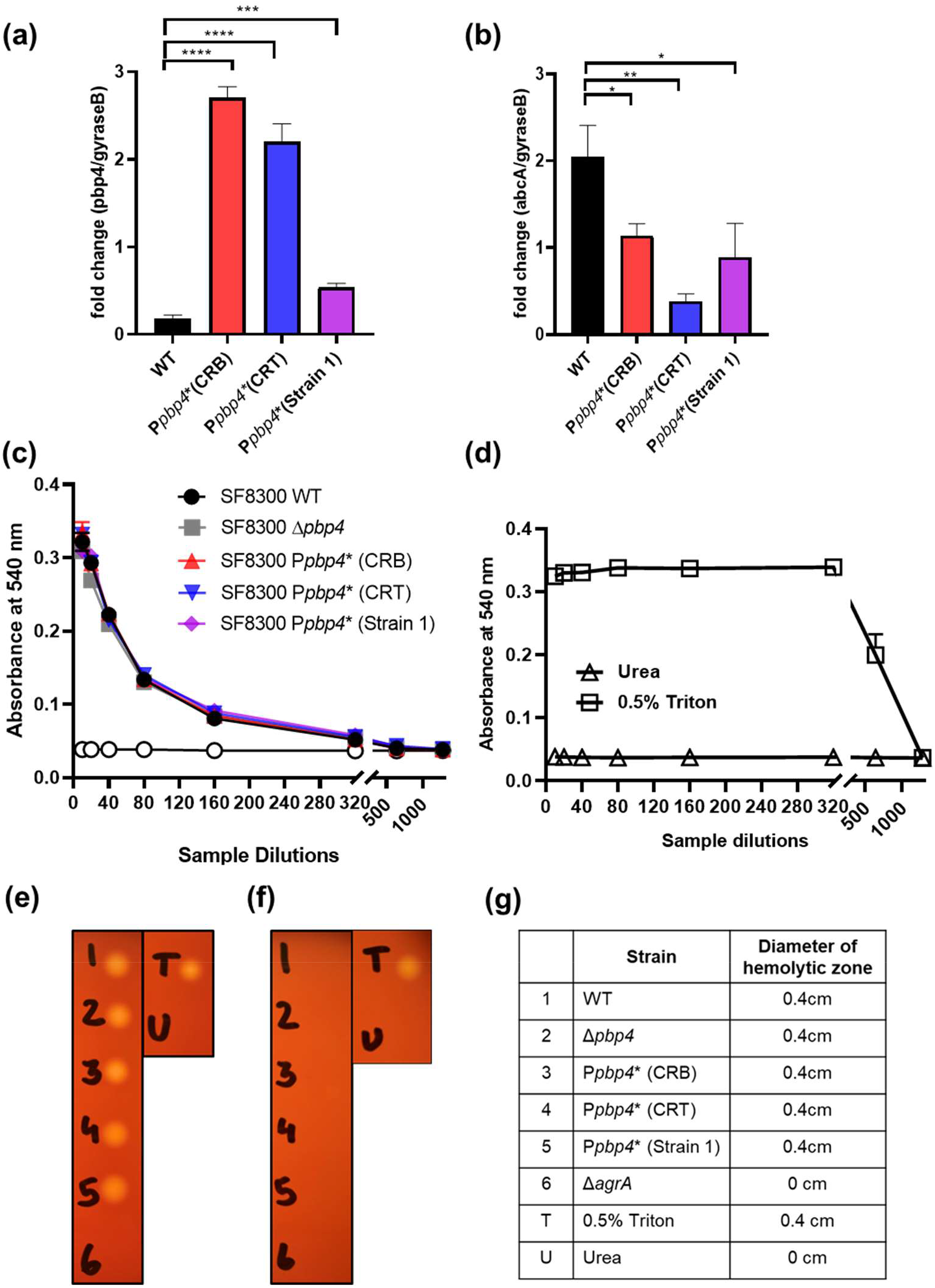
promoter mutations cause an upregulation of *pbp4* and downregulation of *abcA*. **(a) qRTPCR analysis of *pbp4***. *pbp4* transcriptional expression was significantly increased in strains containing promoter-associated mutations as compared to the WT strain. (SF8300 WT versus SF8300 P*pbp4*\* (CRB), P-value < 0.0001, SF8300 WT versus SF8300 P*pbp4*\* (CRT), P-value < 0.0001, SF8300 WT versus SF8300 P*pbp4*\* (Strain 1), P-value = 0.0004) **(b) qRTPCR analysis of *abcA***. *abcA* transcriptional expression was significantly in strains containing promoter-associated mutations as compared to the WT strain. (SF8300 WT versus SF8300 P*pbp4*\* (CRB), P-value = 0.0161, SF8300 WT versus SF8300 P*pbp4*\* (CRT), P-value = 0.0016, SF8300 WT versus SF8300 P*pbp4*\* (Strain 1), P-value = 0.0203) **(c) Hemolysis assay**. Analysis of the ability of butanol extracted Phenol Soluble Modulins (PSMs) to lyse sheep erythrocytes was carried out using a hemolysis assay. There was no difference detected in the hemolytic abilities among the WT strain and strains containing promoter-associated mutations. Δ*agrA* was used as a control, which showed no hemolytic activity. **(d) Controls used for hemolysis assay of sheep erythrocytes**. Hemolysis assay of decreasing concentrations of 8M Urea indicated that urea in itself did not have any hemolytic ability. 0.5% Triton X-100 was used as a positive control, which showed maximum hemolysis. **Analysis of the hemolytic ability of butanol extracted Phenol Soluble Modulins (PSMs) by spotting them onto blood agar-TSA plates**. There was no difference in hemolysis among WT and promoter mutant strains when PSMs were: **(e)** diluted 10 times **(f)** diluted 100 times **(g)** Diameter of the hemolytic zone detected on the blood agar-TSA plates for PSMs from the selected strains.

One of the important functions performed by AbcA is the export of cytolytic toxins such as Phenol Soluble Modulins (PSMs) that can lyse erythrocytes and neutrophils (20). In order to determine the effect of AbcA downregulation and its subsequent effect on its activity, we measured the amount of PSMs exported by WT cells, and cells that contained the promoter-associated mutations. Using 1-Butanol, PSMs were extracted from culture supernatants and re-suspended in urea, following which their ability to lyse sheep erythrocytes was measured by performing hemolysis assays as previously shown (21). It was seen that none of the strains containing promoter-associated mutations differed in their hemolytic capabilities when compared to the WT strain, as they all showed equal amounts of hemolysis at the different dilutions used **(Figure 2c)**. A Δ*agrA* strain was used as a negative control, which showed no hemolysis at all. Urea was used as a control, which by itself did not have any hemolytic abilities, whereas 0.5% Triton X-100 was used as a positive control that showed maximum hemolysis **(Figure 2d)**. Along with using sheep blood for the hemolysis assay, 10x and 100x dilutions of the extracted PSMs were spotted onto blood-TSA plates to analyze differences in hemolysis levels **(Figure 2e-2g)**. The pattern of hemolysis seen on the plates also indicated that there was no difference in the hemolytic abilities between the WT and the mutant strains. The Δ*agrA*, urea and Triton X-100 controls were used here, too. Together, these results confirmed that the β-lactam resistant phenotypes seen with promoter-associated mutations were due to PBP4 overexpression and that AbcA had no role in it.

### *S. aureus* strains with *pbp4* promoter-associated mutations were less virulent to *C. elegans* compared to the wild-type strain

Along with a role in β-lactam resistance, PBP4 has also been associated with pathogenesis. However, its exact role, if any, remains undetermined, as there have been contrasting reports regarding PBP4’s role in pathogenesis (15, 16). We attempted to understand the role of PBP4’s role in pathogenesis, under wild-type and overexpressed conditions using a *C. elegans* infection model. We selected one representative strain containing promoter-associated mutations, namely SF8300 P*pbp4*\* (CRB) and used it to perform infection studies with *C. elegans*. Age synchronized worms were infected with 1.5 × 10^5^ bacteria and were incubated for 3 days, following which they were assessed for worm survival. Worms that responded to mechanical stimulus were considered as live, whereas worms that did not respond were counted as dead. Compared to worms infected with WT cells, where the survival rates for the worms were approximately 30%, worms infected with P*pbp4*\* (CRB) had significantly higher survival rates (55%) (SF8300 WT versus SF8300 P*pbp4*\* (CRB), P-value = 0.0021) **(Figure 3a)** indicating that the presence of promoter-associated mutations led to the decreased killing and thus decreased virulence in *C. elegans*. Infection with SF8300 Δ*pbp4* also resulted in decreased worm survival, similar to the results obtained by infection with WT cells (SF8300 P*pbp4*\* (CRB) versus SF8300 Δ*pbp4*, P-value = 0.052). The *E. coli* strain OP50 was used as a control, where worms displayed 100% survival, indicating that the killing detected was due to S. *aureus* virulence.

**Figure 3:**
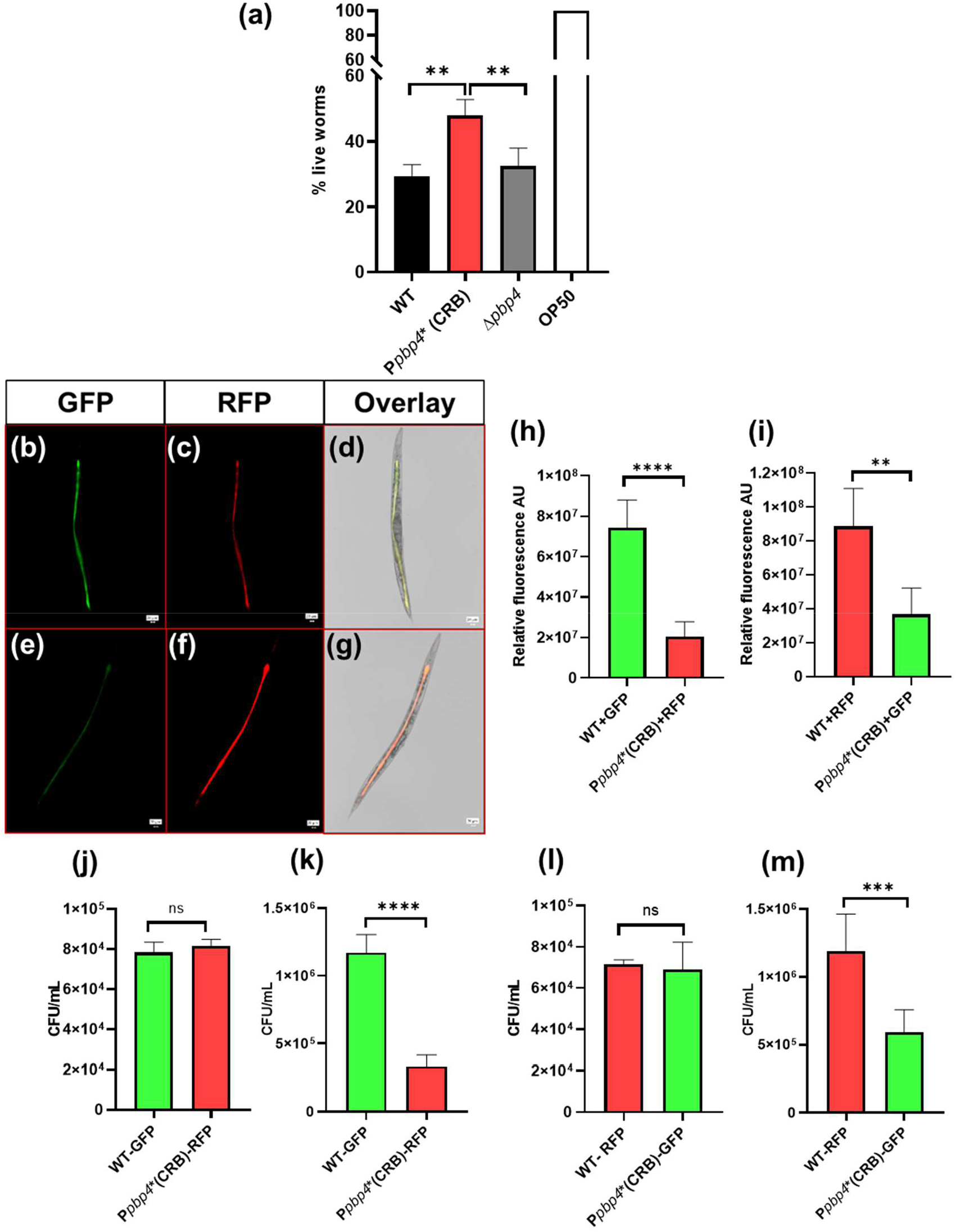
Cells with overexpressed PBP4 have decreased virulence. ***(a) C. elegans* infection assay**. Killing assay with isogenic strains demonstrated that SF8300 WT and SF8300 Δ*pbp4* strains had decreased worm survival (30%) compared to SF8300 P*pbp4*\* (CRB), that had increased worm survival (55%) indicating at decreased *S. aureus* virulence. (SF8300 WT versus SF8300 P*pbp4*\* (CRB), P-value = 0.0021, SF8300 P*pbp4*\* (CRB) versus SF8300 Δ*pbp4*, P-value = 0.052, SF8300 WT versus SF8300 Δ*pbp4*, P value = ns) **Fluorescence microscopy**. Microscopy for *C. elegans* infected with GFP or RFP expressing isogenic strains demonstrated that SF8300 P*pbp4*\* (CRB) had a significantly decreased ability to colonize *C. elegans* compared to WT, after 3 days of infection using a 20X objective lens. Scale bar = 20 µm. **(b)** Infection of *C. elegans* with SF8300 WT + GFP **(c)** Infection of *C. elegans* with SF8300 P*pbp4*\* (CRB) + RFP **(d)** Merged image of *C. elegans* infected with SF8300 WT + GFP and SF8300 P*pbp4*\* (CRB) + RFP **(e)** Infection of *C. elegans* with SF8300 WT + RFP **(f)** Infection of *C. elegans* with SF8300 P*pbp4*\* (CRB) + GFP **(g)** Merged image of *C. elegans* infected with SF8300 WT + RFP and SF8300 P*pbp4*\* (CRB) + GFP. **Analysis of GFP and RFP signals following fluorescent microscopy of *C. elegans* infected with:** **(h)** An equal number of SF8300 WT + GFP and SF8300 P*pbp4*\* (CRB) + RFP demonstrated that there was a significantly increased GFP signal compared to RFP signal from within the worms analyzed (N = 5) indicating that there was increased colonization of the WT strain compared to Ppbp4* (CRB). SF8300 WT + GFP versus SF8300 P*pbp4*\* (CRB) + RFP, P-value < 0.0001. **(i)** Equal number of SF8300 WT + RFP) and SF8300 P*pbp4*\* (CRB) + GFP) demonstrated that there was significantly increased RFP signal compared to GFP signal from within the worms analyzed (N = 5) indicting that there was increased colonization of the WT strain compared to P*pbp4*\* (CRB). SF8300 WT + RFP) versus SF8300 P*pbp4*\* (CRB) + GFP, P-value = 0.0026. **CFU determination**. **(j)** Inoculum on Day 0, before *C. elegans* infection with SF8300 WT + GFP and SF8300 P*pbp4*\* (CRB) + RFP demonstrated that cells of each strain were used in similar proportions. SF8300 WT + GFP versus SF8300 P*pbp4*\* (CRB) + RFP, P-value = ns. **(k)** Plating of bacteria obtained from the gut of *C. elegans* after 3 days of infection with bacteria indicated that there was a higher proportion of GFP expressing SF8300 WT colony forming units (CFU) compared to RFP expressing SF8300 *Ppbp4*\* (CRB) CFUs. SF8300 WT + GFP versus SF8300 P*pbp4*\* (CRB) + RFP, P-value < 0.0001. **(l)** Inoculum on Day 0, before *C. elegans* infection with SF8300 WT + RFP and SF8300 P*pbp4*\* (CRB) + GFP demonstrated that cells of each strain were used in similar proportions. SF8300 WT + RFP versus SF8300 P*pbp4*\* (CRB) + GFP, P-value = ns. **(m)** Plating of bacteria obtained from the gut of *C. elegans* after 3 days of infection with bacteria indicated that there was a higher proportion of RFP expressing SF8300 WT CFUs compared to GFP expressing SF8300 P*pbp4*\* (CRB) CFUs. SF8300 WT + RFP versus SF8300 P*pbp4*\* (CRB) + GFP, P-value = 0.0001.

We carried out further experiments with *C. elegans* infection to determine why the P*pbp4*\* (CRB) strain showed increased survival. SF8300 WT and SF8300 P*pbp4*\* (CRB) strains were introduced with constitutively expressing GFP and RFP respectively, via the constitutively expressing plasmid, *pTX*_*Δ*_, thus generating strains SF8300 WT + GFP and SF8300 P*pbp4*\* (CRB) + RFP. A competition-killing assay was performed, where an equal number of SF8300 WT and SF8300 P*pbp4*\* (CRB) cells were used to infect worms. After 3 days of infection, fluorescence microscopy was carried out where the GFP and RFP signals from within the gut of each worm was was measured. On subsequent analysis of the fluorescent signals, it was observed that there was a significantly increased GFP signal detected, as compared to RFP **(Figure 3b-3d, Figure 3h)**. This indicated that there was a higher proportion of SF8300 WT cells colonized within the gut of *C. elegans*, compared to SF8300 P*pbp4*\* (CRB) cells; (SF8300 WT + GFP versus SF8300 P*pbp4*\* (CRB) + RFP, P-value < 0.0001). The experiment was repeated by interchanging the plasmids containing fluorescent proteins, i.e. with the strains SF8300 WT + RFP and SF8300 P*pbp4*\* (CRB) + GFP **(Figure 3e-3g, Figure 3i)**. Here, an increased RFP signal as compared to GFP signal from within the gut of *C. elegans* was detected, ensuring that it was due to the increased colonization of SF8300 WT, and not due to a result of a potential anomaly of using fluorescent proteins (SF8300 WT + RFP versus SF8300 P*pbp4*\* (CRB) + GFP, P-value = 0.0026).

Before infecting *C. elegans* with bacteria containing fluorescent proteins, the initial inoculum was plated onto tetracycline-containing TSA plates **(Figure 3j)**. After determining that the initial inoculum contained a similar number of each of the bacterial strains (SF8300 WT + GFP versus SF8300 P*pbp4*\* (CRB) + RFP, P-value = ns), *C. elegans* were subjected to lysis after 3 days of infection following which the bacteria accumulated within the gut of the worms were enumerated by performing serial dilutions of the lysate and plating them. There was a significantly higher number of GFP-expressing colonies (representing SF8300 WT) on the plate compared to RFP-expressing colonies (representing SF8300 P*pbp4*\* (CRB)) **(Figure 3k**; SF8300 WT + GFP versus SF8300 P*pbp4*\* (CRB) + RFP, P-value < 0.0001). When plasmids were interchanged, there were increased RFP-expressing colonies (representing SF8300 WT) and decreased GFP-expressing colonies (representing SF8300 P*pbp4*\* (CRB)) **(Figure 3l-m)**. Together, the *C. elegans* experiments indicated that the SF8300 P*pbp4*\* (CRB) was unable to colonize the *C. elegans* gut as well as SF8300 WT, leading to decreased virulence.

## Discussion

MRSA is one of the most prominent agents contributing to the significant global antimicrobial resistance burden today (3). However, in recent years, the presence of β-lactam resistance in *S. aureus* strains without *mecA* have been reported (10, 22, 23). In previous studies, we have demonstrated the ability of PBP4 to mediate high-level β-lactam resistance via protein overexpression due to promoter-associated mutations in non-*mecA* strains (7, 17). In this study, we saw that PBP4 could mediate β-lactam resistance independent of PBP2a. As seen by the growth assays, both PBP2a and PBP4 contributed towardss β-lactam resistance, as cells containing the promoter-associated mutations were able to survive the β-lactam challenge more significantly than strains that only contained PBP2a, i.e. the WT strains. This indicated that when present, PBP4 could supplement the action of PBP2a, causing a further increase in resistance and potentially leads to complications in treatment. Current clinical diagnostic and therapeutic protocols are based on categorization of the infecting strains as MRSA or MSSA as the treatment for the former is more aggressive than the latter (24). However, due to the rise in *pbp4*-associated resistance, it is likely that targeting only PBP2a for diagnosis and treatment protocols may not suffice.

PBP4 is a low-molecular-weight (LMW) PBP in *S. aureus* (25). LMW PBPs in other bacteria such as *E. coli (26), B. subtilis* (27), *L. monocytogenes* (28), *S. pneumonia* (29), etc. have been described to possess carboxypeptidase activity, allowing them to maintain the degree of cell wall crosslinking. PBP4 in *S. aureus* has transpeptidase activity along with carboxypeptidase activity, giving it the ability to perform increased, secondary cell wall cross-linking compared to other bacteria (30). Increased PBP4 expression due to promoter-associated mutations leads to a further increase in cross-linking (17, 31), indicating at PBP4’s propensity towards transpeptidase activity over carboxypeptidase activity. This demonstrates the potential of PBP4 from *S. aureus* to be a powerful player in β-lactam resistance. The importance of PBP4 in β-lactam resistance was further reiterated when we saw that the promoter-associated mutations studied here only led to *pbp4* upregulation, and not *abcA*. AbcA can export various antimicrobial agents, making it an important mediator of antibiotic resistance (14, 19). However, based on the qRTPCR analysis and hemolysis assay, it was clear that AbcA did not play any role in the resistant phenotype associated with the promoter mutations, and that it was all attributed to PBP4 overexpression. It is likely that the presence of mutations possibly led to altered binding of transcriptional factors to the regulatory region, leading to overexpression of PBP4. Reports have suggested that *pbp4* and *abcA* regulation are independent of each other (14), however, since the studied mutations also caused the downregulation of *abcA*, it appears that the regulation of both of these genes are at least partially interdependent, and they potentially share some of their regulatory sequences as well as some regulatory factors. Along with AbcA, *S. aureus* also has another known exporter of PSMs, namely the Pmt system (21). It is likely that Pmt compensated for the export of PSMs during *abcA* downregulation, allowing for similar levels of hemolytic activities between various strains.

In *S. aureus*, PBP4 maintains the level of peptidoglycan cross-linking by its transpeptidase and carboxypeptidase properties, thus striking a balance between appropriate cross-linking of peptidoglycan monomers and allowing for anchoring of surface proteins that are necessary for *S. aureus* virulence (32). Our previous studies demonstrated that this well-balanced cell wall dynamic was disrupted in the form of increased cell wall cross-linking due to overexpression of PBP4 (7, 31). However, the effect of this disruption on the abundance of surface-anchored proteins is unknown. As these surface proteins play vital roles in host attachment, colonization, and infection, we examined the virulence of a strain containing promoter-associated mutations using *C. elegans*, as this strain exhibited increased PBP4 expression. *C. elegans* has proven to be a useful infection model system to study host-pathogen interactions previously and have given multiple clues about host response to pathogenic bacteria (33). *S. aureus* cells mediate *C. elegans* killing by accumulating with the gut of the worm, leading to intestinal distension and subsequent worm death (34). Based on our results, strains with overexpressed PBP4 were unable to colonize within the gut of the worms to a sufficient enough degree to cause high levels of accumulation and worm death, compared to the WT cells. Gut epithelial cells of *C. elegans* have been described to have similarities with human intestinal epithelial cells, suggesting the possibility that strains containing increased PBP4 expression may have decreased ability to colonize humans, as *S. aureus* has been shown to also colonize and infect human intestinal cells, making this study biologically relevant (35). It is likely that the inability of P*pbp4*\* (CRB) cells to colonize was due to a decrease in surface-anchored proteins due to increased PBP4-mediated cross-linking of the peptidoglycan. In *S. aureus*, repeating monomers of the peptidoglycan consisting of penta-peptide stems are cross-linked via a penta-glycine bridge (36). The terminal glycine of this pentaglycine bridge forms a peptide bond with the D-alanine of the penta-peptide stem, via PBP4’s transpeptidase activity (37). However, this glycine is also the site at which sortase A mediated anchoring of cell surface proteins takes place (38). Due to PBP4-mediated increase of cell wall cross-linking, we hypothesize that there aren’t sufficient free glycine sites available for the sortase A mediated attachment of surface proteins, thus leading to decreased virulence **(Figure 4)**. Further studies are required to understand the biology that leads to the decreased virulence phenotype of *S. aureus* strains having *pbp4*-promoter associated mutations, which can potentially help in exploiting this phenomenon for treatment purposes.

**Figure 4:**
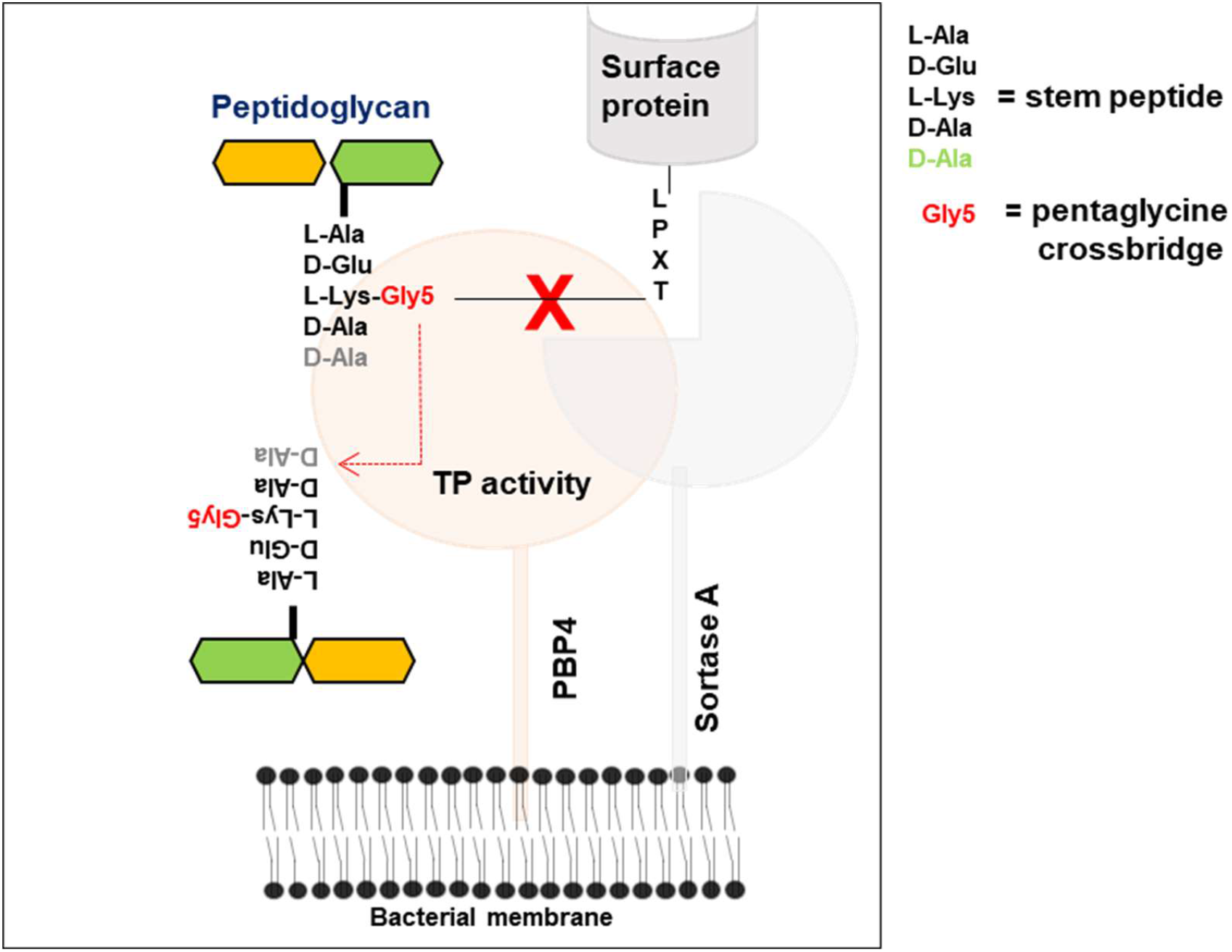
Increased cell-wall cross-linking due to PBP4 overexpression can result in decreased anchoring of sortase A mediated cell surface associated proteins.

## Material and Methods

### Bacterial strains and plasmids

*S. aureus* strains were all cultured at 37°C in TSB (Tryptic Soy Broth), with agitation at 180 rpm. Promoter-associated mutations in *S. aureus* were introduced via Splice-Overlap PCR and allelic replacement as described previously (39), using the plasmid pJB38 (40). The GFP and RFP encoding regions were amplified from the integration plasmids pGFP-F and pRFP-F respectively (40), and were cloned into the constitutively expressing plasmid *pTX*_*Δ*_ as described previously (17). The plasmid was introduced into RN4420 by electroporation, following which they were introduced in SF8300 or SF8300 P*pbp4*\* (CRB) via phage transduction using Φ11. All strains, primers and plasmids used in this study are listed in **Tables 1, 2 and 3**, respectively.

**Table 1:**
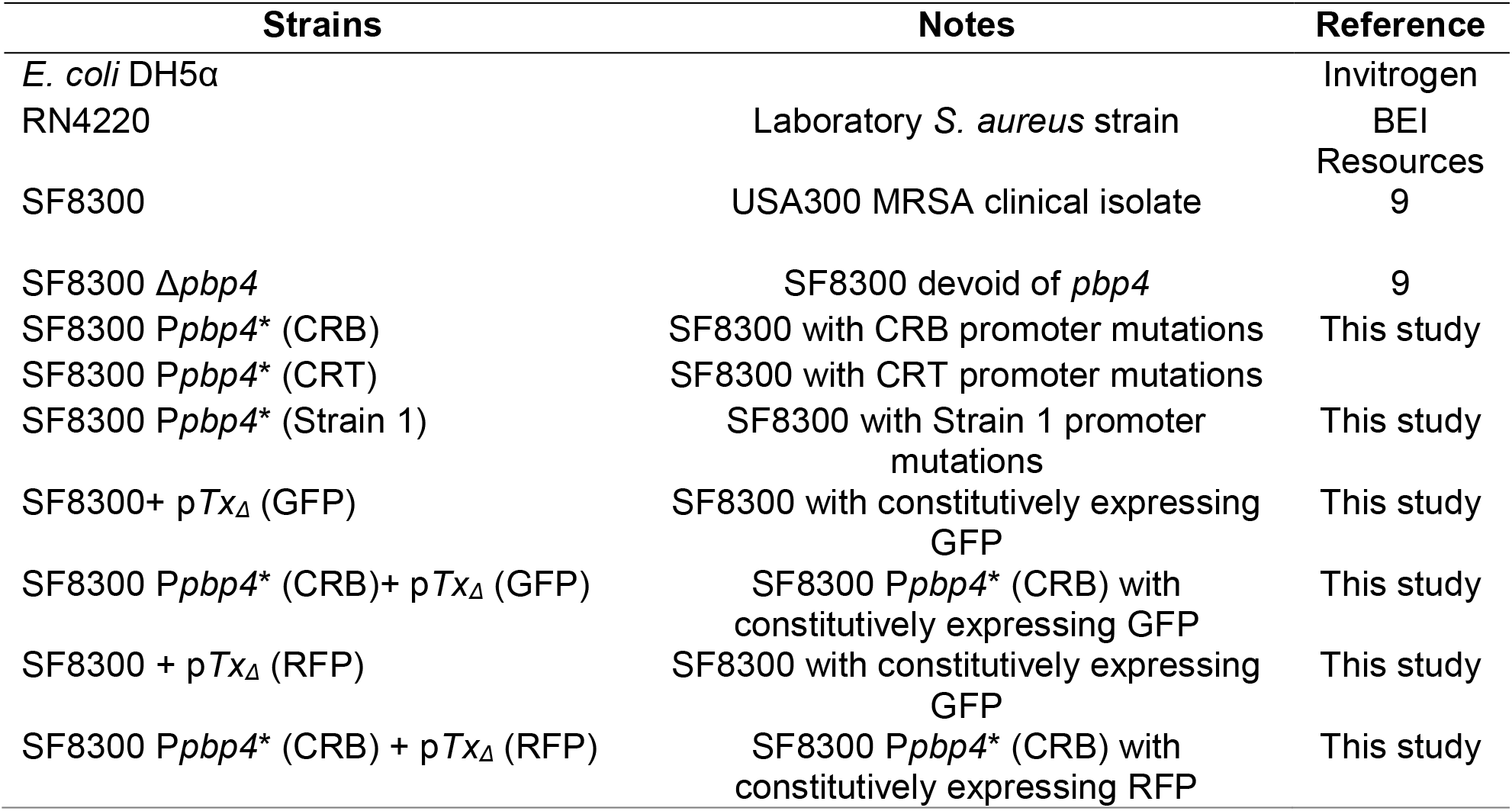
Strains used in this study.

**Table 2:**
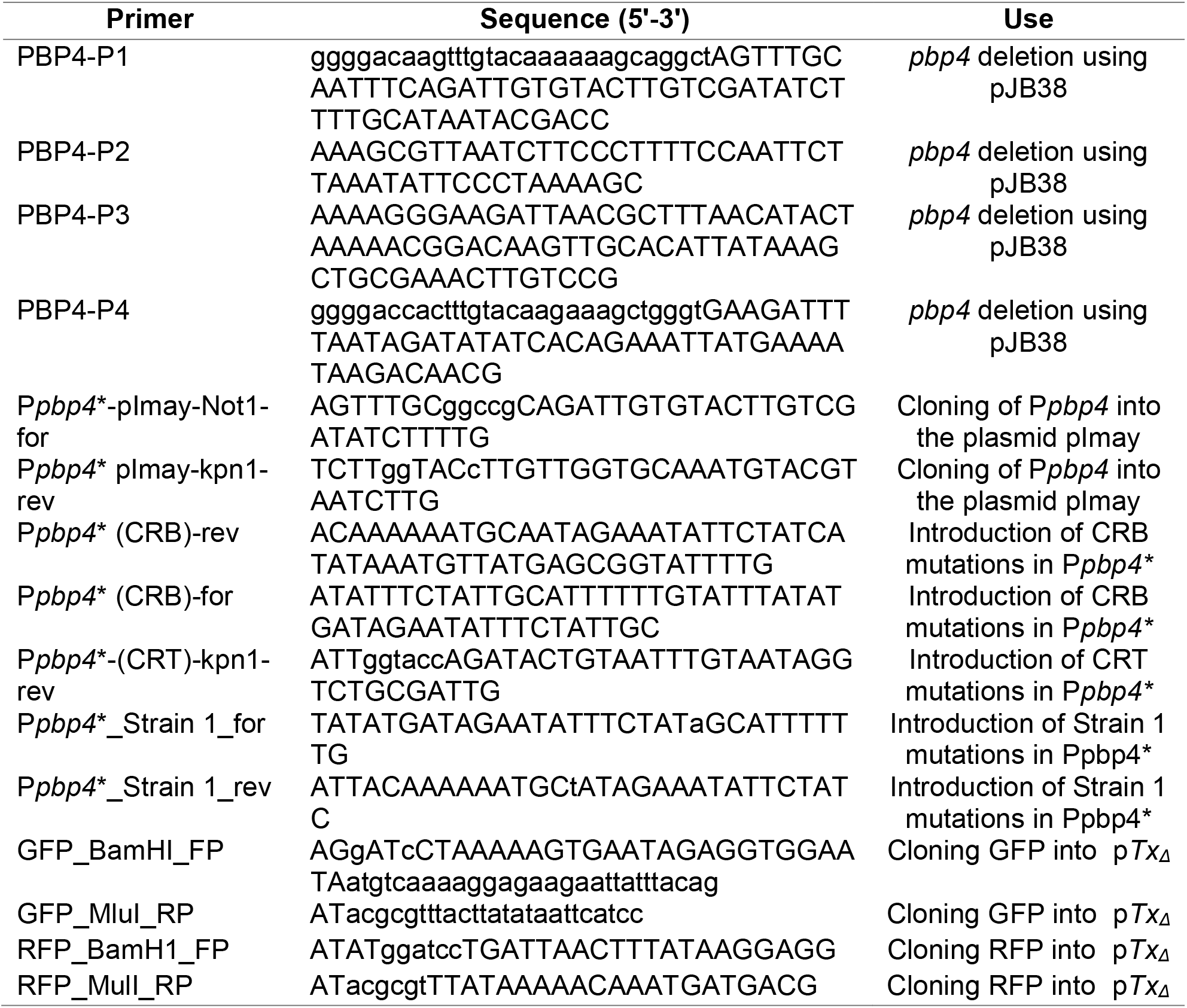
Primers used in this study.

**Table 3:** Plasmids used in this study.

| Plasmid | Notes | Reference |
| --- | --- | --- |
| pImay + *Ppbp4* (CRB) | Insertion of *Ppbp4* promoter mutation | This study |
| pJB38 + *Ppbp4* (CRT) | Insertion of *Ppbp4* promoter mutation | This study |
| pJB38 + *Ppbp4* (Strain 1) | Insertion of *Ppbp4* promoter mutation | This study |
| pJB38 + $\Delta pbp4$ | Deletion of *pbp4* | |
| p*Tx<sub>Δ</sub>* + GFP | Constitutive expression of GFP | This study |
| p*Tx<sub>Δ</sub>* + RFP | Constitutive expression of RFP | This study |

### Growth curve assays

Growth assays were performed using the automated microbiology growth curve analysis system, Bioscreen C (Growth Curves USA). Overnight cultures of bacteria were diluted to OD = 0.1 in TSB with or without antibiotics, and 200 µL was added to each well of a honeycomb bioscreen C plates in triplicates. The assay was carried out for 20 hours with continuous orbital shaking at 37°C. Each condition was in triplicates, and the experiment was performed twice to ensure reproducibility.

### Immunoblotting

Overnight cultures of bacteria were subcultured into 50mL flasks containing TSB such that the initial OD_600_ of the flasks was 0.1. The cells were cultured to OD_600_ = 1, following which cells were collected and resuspended in PBS containing CompleteMini protease inhibitor cocktail (Roche). The cells were mechanically lysed using the FastPrep (MP Biochemicals) and whole cell lysates were obtained. The cell membrane fraction was isolated from the lysates by performing ultracentrifugation at 66000 g for 1 hour (Sorvall WX Ultra 80 Centrifuge, Thermo Fisher Scientific). After resuspending the obtained pellet with PBS, protein estimation was carried out using the Pierce BCA Protein Assay kit (Thermo Fisher). The samples were separated by performing SDS-PAGE on a 10% gel, following which they were transferred onto a low-fluorescence PVDF membrane (Millipore). Blocking was performed for 1 hour (5% skimmed milk in Tris buffered saline containing 0.5% Tween), primary antibody staining was carried out overnight at 4C (polyclonal anti-PBP4, custom antibody from Thermo Fisher, 1:1000) and secondary antibody staining was performed using an anti-rabbit antibody (Azure anti-rabbit NIR700, 1:20000 dilution). The blots were imaged using the Azure C600 imager and analysis was performed using ImageJ.

### Qrtpcr

Overnight cultures of bacteria were subcultured into 50mL flasks containing TSB such that the initial OD_600_ of the flasks was 0.1, and cells were allowed to grow for 4 hours, at which point 5 × 10^9^ bacterial cells were harvested and washed. Cells were lysed using FastPrep (MP Biochemicals) following which RNA isolation was performed using the Qiagen RNeasy Mini kit. On confirmation of RNA quality, cDNA synthesis was performed using the SuperScript® IV Reverse Transcriptase (RT) kit. qRT-PCR was performed using SYBR PCR Mastermix in an ABI 7500 system (Applied Biosystems) primers for *pbp4, abcA*, and the housekeeping gene, *gyrB*.

### Butanol Extraction of PSMs

PSMs were extracted from culture supernatants as previously described (21, 41). Briefly, overnight cultures of bacteria were subcultured into 50mL flasks containing TSB such that the initial OD_600_ of the flasks was 0.1. After 24 hours, the cells were collected, centrifuged for 30 minutes, and 30mL of supernatant from each strain was collected to which 10 mL 1-Butanol was added. The samples were mixed by shaking at 180 rpm for 2 hours at 37C. Samples were then centrifuged, and 7 mL of the upper layer from each sample was collected. The samples were dried using a vacuum centrifuge (Eppendorf) and the dried pellet was re-suspended in 8M urea. Samples were diluted 10-fold for hemolysis experiments.

### Hemolysis assay

2% sheep blood was prepared using chilled PBS and was washed twice to get rid of lysed erythrocytes. After washing, 100 µL of the blood was added to a 96 well round bottom plate, to which 100 µL of samples of PSMs extracted with butanol was added in decreasing concentrations (from 1/10^th^ to 1/1280^th^ of the sample). The plates were incubated for 1 hour at 37°C, following which they were centrifuged at 1500 rpm for 10 minutes. 100 µL of the supernatant was collected and transferred onto a flat bottom 96 well plate. Absorbance was measured at 540 nm using SpectraMax M5 (Molecular Devices). 0.5% Triton was used as a control.

2 µL of butanol extracted PSMs was spotted onto TSA-blood plates (5% sheep blood) for each sample. Urea was spotted as a negative control and 0.5% triton was spotted as a positive control. On drying of the samples, plates were incubated at 37°C overnight.

### *C. elegans* – *S. aureus* infection assays

*C. elegans* were obtained from the Caenorhabditis Genetics Center (CGC). Liquid killing assay was performed as previously described (42), with certain modifications. *C. elegans* strain used in this study was DH26 (rrf-3(b26) II), a temperature-sensitive, spermatogenesis defective strain. Briefly, adult hermaphrodites were age-synchronized (43) using household bleach and 5N NaOH and allowed to grow on NGM plates for 40 hours at 26°C. 15 young adult worms were picked from this plate and added to 96 well plates containing 100 µL liquid killing media (80% M9 buffer, 20% TSB, 100 µg/mL cholesterol, and 7.5 µg/mL Nalidixic acid) and 1.5 × 10^5^ *S aureus* cells. Wells containing OP50 in media (80% M9 buffer, 20% LB and 200 µg/mL cholesterol) were included as controls. The plate was incubated at 26°C for 3 days following which live and dead worms were counted. The experiment was performed thrice, in triplicates in order to ensure reproducibility.

For the competition assay, WT + GFP and P*pbp4*\* (CRB) + RFP cells were mixed in equal amounts before adding 1.5 × 10^5^ cells to each well. The media also contained 12.5 µg/mL tetracycline, which is the selection antibiotic for the *pTX*_*Δ*_ plasmid.

### Fluorescence microscopy and Image analysis

Fluorescence microscopy and analysis were performed as previously described (44). After 3 days of infection with *S. aureus*, worms were transferred from the wells to 1.5 mL microcentrifuge tubes. The worms were washed with M9 buffer containing 10 mM sodium azide, following which they were treated with 100 µg/mL gentamicin for 30 minutes, twice. Following the gentamicin treatment, worms were washed thrice with M9 buffer containing 10 mM sodium azide. The worms were then placed onto an agar pad (2% agar) on glass slides, and fluorescence microscopy was performed using the Keyence BZ-X800 All-in-one Fluorescence microscope using the 20X objective. The exposure time and all other settings remained the same for all samples.

The BZX Analyzer software was used to measure fluorescence intensities within a worm. After stitching images wherever necessary, the “Hybrid cell count” function was used to select the area of the worm and measure fluorescence intensities for GFP and RFP. For each condition, at least 5 worms were imaged and analyzed.

### Gut CFU determination

Gut bacteria enumeration was performed as previously described (45). After the gentamicin treatment and wash steps as described above, the worms were resuspended in 250 µL M9 buffer containing 10 mM sodium azide, mixed with 200 mg 1mm Zirconia beads (Research Products International), and lysed by vortexing the tubes for 2 minutes. Before lysis, 20 µL of the supernatant was collected for plating. Following lysis, samples were diluted to 10^−4^ and plated onto TSA plates containing tetracycline, and incubated at 37°C. The next day, colonies were counted based on the color they expressed. The experiment was performed twice, in triplicates to ensure reproducibility. Each tube contained 10 worms, and the experiment was performed twice, in triplicates.

### Sequencing

Fidelity of all the mutants and plasmid constructs was validated through Sanger sequencing (Eurofins Genomics, USA).

### Bioinformatics and statistical analysis

Statistical analyses were performed using GraphPad Prism. Comparisons between groups were analyzed by two-tailed Student’s t-test whenever stated. DNA sequence analysis was performed using DNAstar software.

## Acknowledgments

The authors are thankful to Dr. Jenniffer Powell, Gettysburg College, for her assistance and guidance in setting up the *C. elegans* infection model system in the Chatterjee Lab. *C. elegans* strains were obtained from the Caenorhabditis Genetics Center (CGC), which is funded by NIH Office of Research Infrastructure Programs (P40 OD010440). This work was funded by NIH grant 2R01AI100291 provided to Som S. Chatterjee.

